# A unified eIF1A^+^ luminal cells-centered hypoxic and “cold” tumor microenvironment promotes PCa progression among different subtypes

**DOI:** 10.1101/2025.04.17.649316

**Authors:** Lilin Wan, Enyao Huang, Zicheng Xu, Bing Zheng, Xuefei Ding, Chunyan Chu, Tiange Wu, Shengrong Chen, Junyong Zhuang, Ming Chen, Rong Na, Shenghong Ju, Dingxiao Zhang, Wenchao Li, Yifei Cheng, Bin Xu

## Abstract

PCa is a malignancy with high heterogeneity arises from both tumor microenvironment (TME) and histological subtypes. To achieve superior clinical efficacy, it is imperative to identify unified progression drivers within such heterogeneity in PCa, thereby enabling the development of more consistent and effective therapeutic strategies. In this study, we applied imaging mass cytometry to stain 39 proteins on 71 tissues comprising para-cancer, low-grade acinar adenocarcinoma (LgPAC), high-grade PAC (HgPAC), intraductal carcinoma (IDC) and ductal adenocarcinoma (DAC) tissues, obtaining the spatial proteomic landscape of 345,233 single cells. We discovered a hypoxic eIF1A^+^ luminal epithelial (LE) cluster exhibiting both translational activation and immunosuppressive properties, which was enriched in high-risk PCa including HgPAC, IDC and DAC, and associated which tumor progression and unfavorable prognosis. Additionally, eIF1A^+^ LE orchestrates the formation of an immunologically “cold” TME, characterized by the low infiltration of anti-cancer immune cells including PD1^-^ T cells CD163^-^ macrophages. Through *in vivo* validation and investigator-initiated study, we further found that Homoharringtonine, an inhibitor targeting protein translation process, effectively inhibited PCa growth, alleviated hypoxic microenvironment, and enhanced immune cell infiltration. This study is the first to apply spatial proteomics to delineate consistent molecular features across histological subtypes, providing transformative insights for clinical management of PCa.

## Introduction

Prostate cancer (PCa) is currently the most frequent diagnosed malignancy and the second most common factor in cancer-related deaths in men worldwide ^1^. Since the establishment of androgen deprivation therapy (ADT) in 1941, it is the standard treatment for PCa. However, after 18-24 months ADT, the non-metastatic PCa patients will develop castration-resistant PCa (CRPC) ^2,3^. Though novel hormonal therapies and chemotherapy have delayed clinical progression to some degree, treatment options remain limited for advanced-stage PCa. Immune checkpoint inhibitors (ICIs) have significant efficacy in some tumors, but due to the immunosuppressive characteristics of tumor microenvironment (TME) (i.e., absence of anti-tumor T-lymphocytes and infiltration of macrophages ^4,5^), the ICIs efficacy is unsatisfactory in PCa ^6,7^. Hence, there is an urgent necessity to explore new therapeutic directions for advanced PCa.

PCa is a malignancy with high heterogeneity, which arises from both TME and histological subtypes. To achieve superior clinical efficacy, it is imperative to identify unified progression drivers within such heterogeneity in PCa, thereby enabling the development of more consistent and effective therapeutic strategies. PCa histological subtypes include acinar adenocarcinoma (PAC), intraductal carcinoma (IDC), ductal adenocarcinoma (DAC) and so on, which represent different clinical characteristics. The IDC is usually associated with higher Gleason scores, larger tumor volumes, neuroendocrine differentiation, and faster disease progression ^8–10^, and is less responsive to ADT, ultimately leading to lower recurrence-free and overall survival rates^11–13^. DAC is a subtype frequently associated with biochemical recurrence and distant metastasis ^14–17^. Unfortunately, there are no systematic studies focusing on the molecular characteristics of different histological subtypes due to the rareness of DAC and IDC types, resulting in insufficient evidence for corresponding treatment approaches. Moreover, technical limitations in isolating relevant tissues from PCa specimens have restricted characterization of rare subtypes (IDC and DAC) to *in situ* analyses, which is an unresolved challenge in the field. Hence, understanding the spatial pattern of distinct histological entities in advanced PCa is crucial to reveal the common underlying progression mechanisms, thus providing insights for new therapeutic strategies.

Imaging mass cytometry (IMC) is a tissue section-based method for *in situ* cellular analysis using metal-labeled antibodies, providing information of cellular proteomic phenotypes and spatial distribution features at a single-cell resolution ^18^. In this study, by applying IMC in para-cancer tissues (PCT) and PCa tissues covering different histological subtypes, we discovered a hypoxic eukaryotic initiation factor 1A (eIF1A)^+^ luminal epithelial (LE) cluster exhibiting both translational activation and immunosuppressive properties, which orchestrates the formation of an immunologically “cold” TME. This eIF1A^+^ LE was enriched in high-risk PCa including high-grade PAC (Hg-PAC), IDC and DAC, and associated which unfavorable prognosis. We also suggested that Homoharringtonine (HHT), an inhibitor targeting protein translation process, effectively inhibited *in vivo* tumor growth, alleviated hypoxic microenvironment, and enhanced immune cell infiltration. This study is the first to apply spatial proteomics to delineate consistent molecular features across PCa histological subtypes, providing transformative insights for clinical management.

## Results

### IMC analysis of para-cancer prostatic and PCa tissues

We first investigated the heterogeneity of the proteomic microenvironment in prostatic tissues through IMC. A tissue microarray (TMA) was generated from 25 para-cancer tissues (PCT), 17 low-grade PAC (LgPAC; PAC with Gleason score 3+3), 7 HgPAC (PAC with Gleason score ≥ 3+4), 11 IDC and 11 DAC, with at least follow-up until April 2025 (**Supplementary Table 1**). In this study, we assigned HgPAC, IDC and DAC to a high-risk while LgPAC to a low-risk PCa group. The histological type of each core on the TMA was determined through a combination of hematoxylin-eosin (H&E) staining and immunohistochemical (IHC) staining for CKH and p63 (**Supplementary Figure 1a**).

The TMA was stained with 39 well-designed metal-labeled antibodies, including established cell type markers and key functional markers (**Supplementary Figure 1b-c**). We then performed IMC analysis, and quantified marker expression in the resulting highly multiplexed images at single-cell resolution, which generated a total of 345,233 cells and formed the “IMC cohort” (**Figure 1a**). The marker panCK was used to identify 156,404 epithelial cells, and classified them into basal epithelia (BE, CK5^hi^) and luminal epithelia (LE, CK5^lo^). The other microenvironment cells included 14,995 T cells (CD3), 2,548 B cells, 4,670 CD163^-^ macrophages, 12,565 CD163^+^ macrophages, 8,767 other antigen presentation cells (APCs), 4,060 other immune cells, 23,537 endothelial cells, 50,966 fibroblasts, 20,851 myofibroblasts, 18,089 smooth muscle cells (SMCs) and 27,781 other stromal cells (**Figure 1b-d** and **Supplementary Figure 1d**). LE constituted the predominant component within the TAM, accounting for 38.5% of the total population (**Figure 1c**).

**Figure 1.**
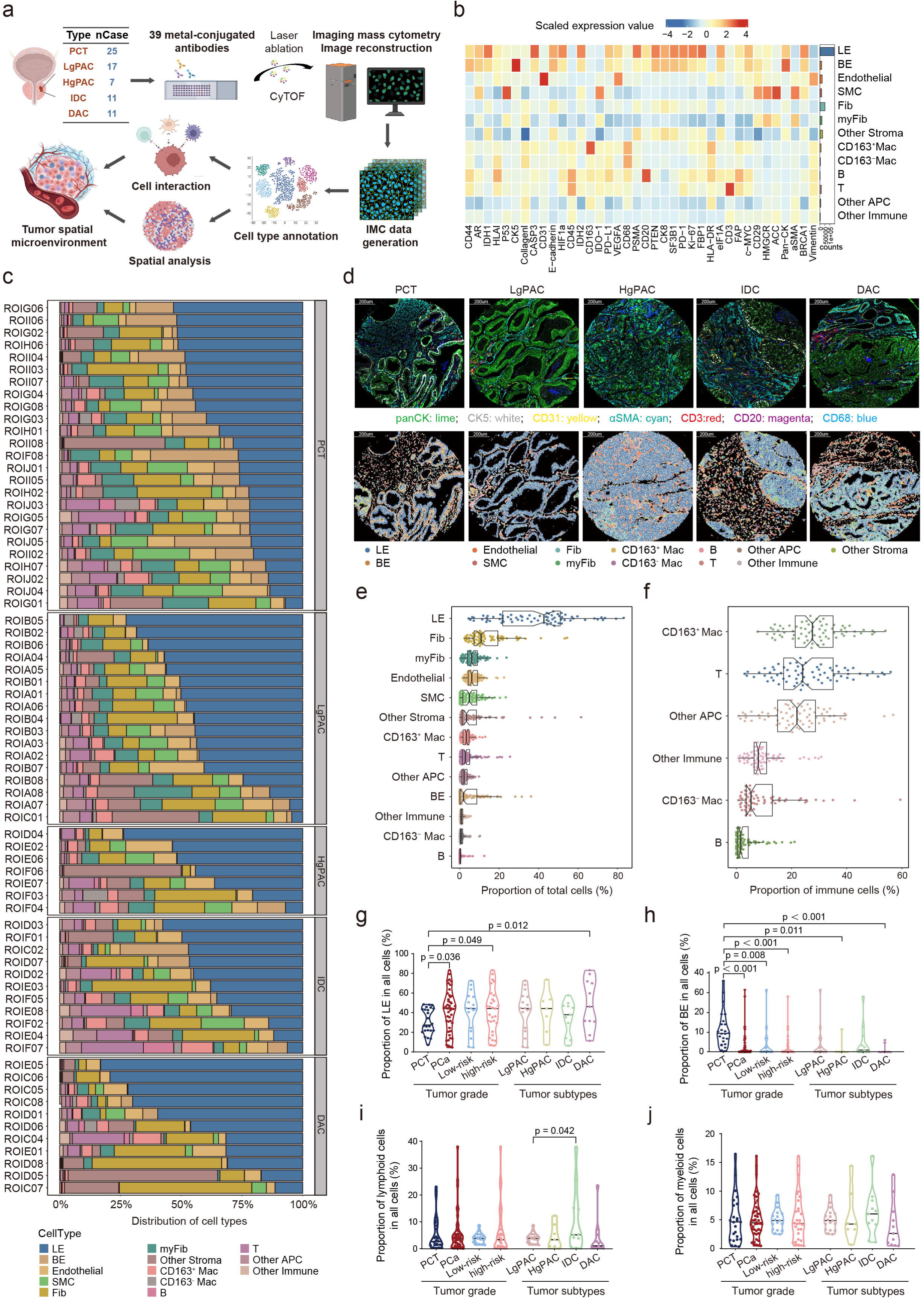
IMC analysis of para-cancer prostatic and PCa tissues. **(a)** The flowchart of IMC analysis. **(b)** Scaled protein expression levels in different cell types. **(c)** The distribution of different cell types among 71 TMA cores. **(d)** Representative IMC images showing the staining of cell type-specific proteins and the distribution of different types of cells in PCT, LgPAC, HgPAC, IDC and DAC. **(e-f)** Boxplots showing the proportion of each type of cells (e) and immune cells (f). **(g-j)** Violin plots comparing the proportion of LE (g), BE (h), myeloid cells (i) and T cells (j) between different types of tissues (PCT, PCa, low-risk PCa, high-risk PCa, LgPAC, HgPAC, IDC and DAC).

We observed significant enrichment of LE in PCa cores, while BE were predominantly enriched in PCT, consistent with pathological features of PCa (**Figure 1g-h**). However, the proportions of LE and BE showed no significant association with different grades or subtypes of PCa (**Figure 1g-h**). Furthermore, across PCT and PCa cores with varying grades or histological subtypes, no significant differences were detected in the proportions of lymphocytes and myeloid cells, except for notably higher lymphocyte infiltration in IDC compared to LgPAC (**Figure 1i-j**). In summary, the abundance of LE is insufficient to predict progression in PCa, highlighting the necessity for further annotations to delineate more precise functional subtypes at proteomic level. On the other hand, as a type of “cold” tumor, the differences in the immune microenvironment among PCa subtypes may be mediated by spatial heterogeneity in cellular distribution rather than variations in infiltration levels.

### eIF1A^+^ LE is associated with progression and poor prognosis of PCa

To identify crucial epithelial subtypes, we assessed the relationship between the expression of key functional markers in epithelia and clinical features. We found that compared to well-established PCa pro-cancer factors, such as AR, PSMA and BRCA1 proteins, eIF1A was significantly associated with more and the most progression linked features (including more tumor quantification, greater pathologic grade, progressed stage and periphery invasion), advanced histological types (HgPAC, IDC and DAC), poorer overall survival (OS) and progression-free interval (PFI) in PCa (**Figure 2a-c**). The TCGA database also showed that *EIF1A* (the key gene encoding eIF1A) expression was significantly higher in PCa than PCT, and higher in Gleason score 8-10 (**Figure 2d-e**). Moreover, *EIF1A* also suggested poorer overall survival in TCGA database (**Supplementary Figure 2a**). In contrast, the previously identified PCa-associated genes, including *AR*, *FOLH1* (the gene encoding PSMA), and *BRCA1*, showed no prognostic value in TCGA database, suggesting that eIF1A is a promising target that may participate in the facilitation of PCa progression (**Supplementary Figure 2b**). eIF1A participates in the scanning process of translation initiation and is a key rate-limiting step in protein translation that ensures protein synthesis accuracy. We further categorized the epithelia into eIF1A^+^ LE (CK5^lo^/eIF1A^hi^), eIF1A^-^ LE (CK5^lo^/eIF1A^lo^), and BE (CK5^hi^) by the expression of CK5 and eIF1A. We found notably enrichment of eIF1A^+^ LE in the tumors, especially in HgPAC, IDC and DAC (**Figure 2f** and **Supplementary Figure 2c-d**). Expectedly, the higher percentage of eIF1A^+^ LE was associated with poorer OS and PFI (**Figure 2g-h**). We then constructed a “tissue validation cohort” consisting of 12 additional cases of PCa to validate the above IMC results, in which 8 cases were histologically confirmed to contain IDC, 5 cases contained DAC, and all possessed PAC component. The results of multiplex immunofluorescence (mIF) staining indicated that CK8 and eIF1A exhibited more pronounced co-localization in HgPAC, IDC, and DAC compared to LgPAC (**Figure 2i**). Further quantitative comparative analysis revealed that both eIF1A expression and eIF1A^+^ LE abundance were elevated in high-risk subtypes (**Figure 2j-k**). All these findings suggested that eIF1A may play an important role in the progression of PCa, and this phenomenon is consistent in different histological subtypes including PAC, IDC and DAC.

**Figure 2.**
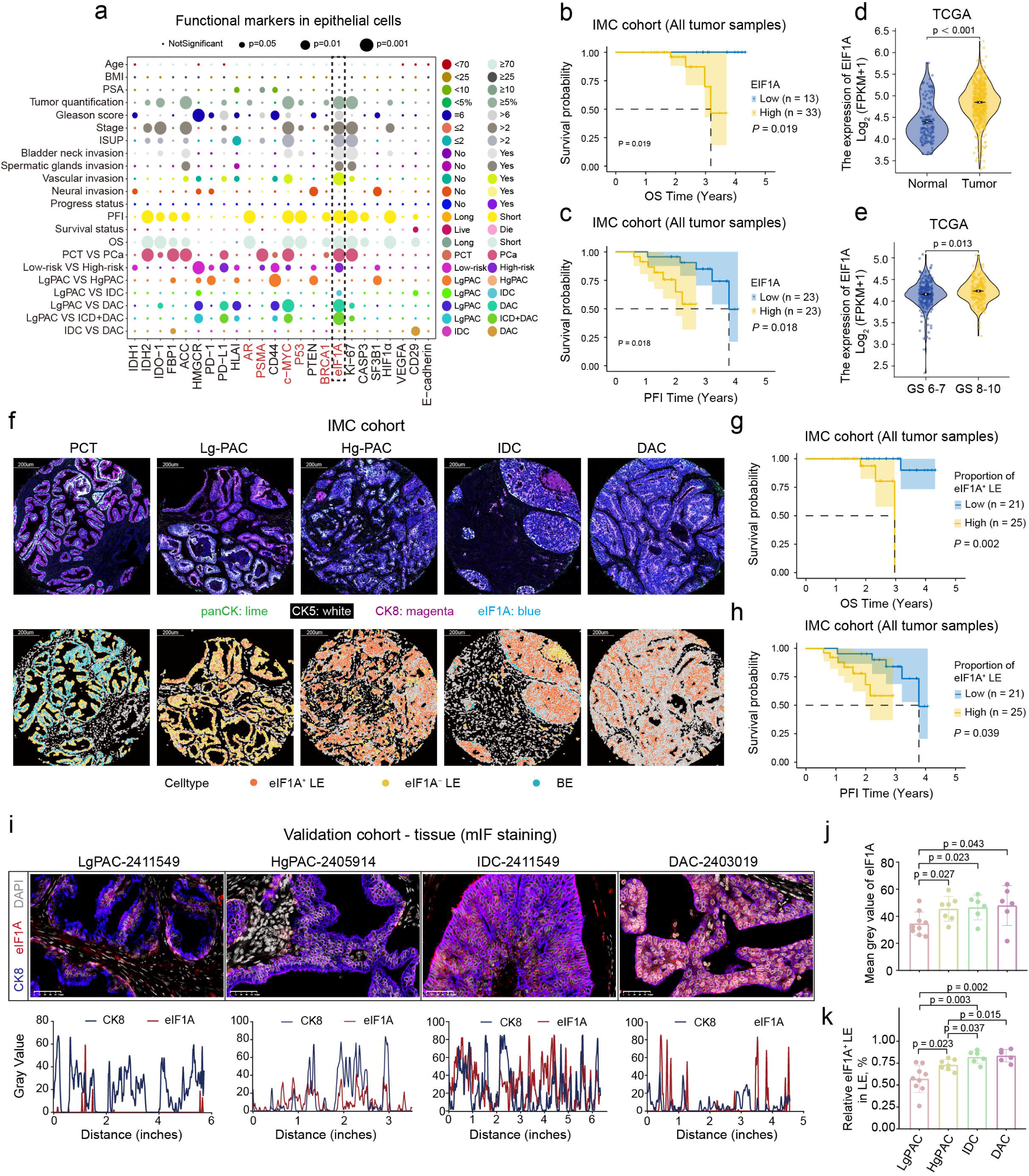
eIF1A^+^ LE is associated with progression and poor prognosis of PCa. **(a)** Dot plot showing the association between the functional markers expression levels of epithelial cells with clinical features. **(b-c)** K-M plots showing the OS (b) and PFI (c) differences between eIF1A-high and -low patients in the IMC cohort. **(d-e)** Violin plots comparing the expression level of *EIF1A* between tumor and normal tissues (d), as well as between Gleason score 8-10 and 6-7 tissues (e) in TCGA-PRAD database. **(f)** Representative IMC images showing the staining of panCK, CK5, CK8 and eIF1A (upper), as well as the distribution of different types of epithelial cells (lower) in PCT, LgPAC, HgPAC, IDC and DAC. **(g-h)** K-M plots showing the OS (g) and PFI (h) differences between patients with high and low proportion of eIF1A^+^ LE in the IMC cohort. **(i)** Representative mIF staining images (upper) and gray value distribution plots (lower) showing the co-localization of CK8 and eIF1A in different types of PCa. **(j-k)** Bar plots comparing the mean gray value of eIF1A staining (j) and the proportion of eIF1A^+^ LE in LE (k) between different types of PCa.

### eIF1A^+^ LE exhibits a hypoxic phenotype

To deeply investigate the characteristics of eIF1A^+^ LE, we compared the expression level of functional markers among epithelial sub types. The results demonstrated that, in addition to the aforementioned PCa-associated proteins, eIF1A+ LE exhibited a distinct hypoxic (HIF-1α^hi^) and metabolically reprogrammed microenvironment. Hypoxia is a hallmark of tumor progression, capable of inducing angiogenesis as well as alterations in glucose, lipid, and energy metabolism. Consistently, we found that eIF1A^+^ LE simultaneously overexpressed VEGFA and CD29, indicating upregulated angiogenesis. Additionally, key rate-limiting enzymes involved in fatty acid synthesis (ACC^hi^), cholesterol synthesis (HMGCR^hi^), and gluconeogenesis (FBP1^hi^) were significantly upregulated. Elevated expression of IDH1 and IDH2 further suggested enhanced tumor energy metabolism (**Figure 3a**). Thus, we further focused on the hypoxic characteristics and the HIF-1α expression of eIF1A^+^ LE. Expectedly, in eIF1A+ LE, HIF-1α was significantly overexpressed in high-than in low-risk PCa (**Figure 3b**).

**Figure 3.**
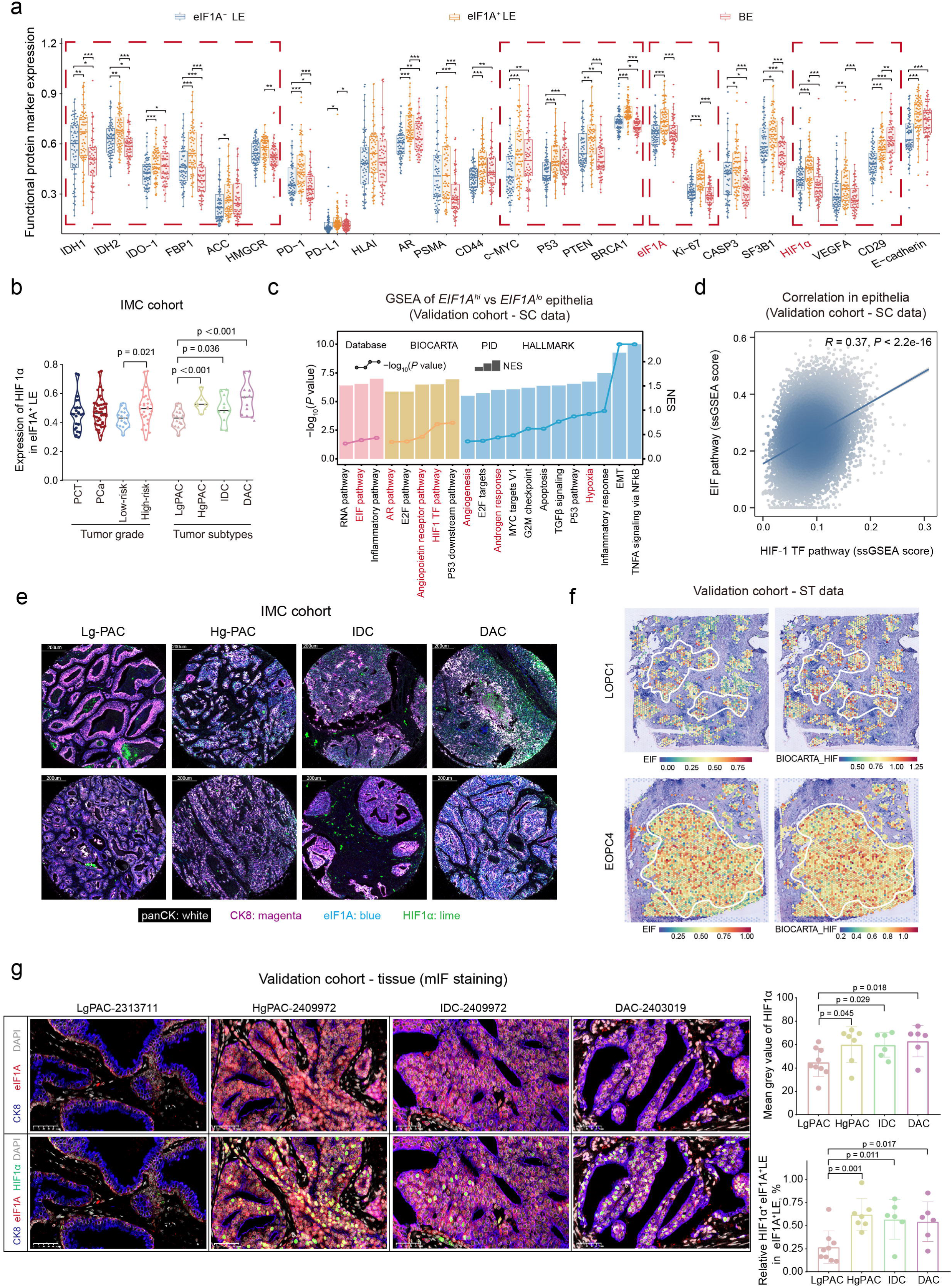
eIF1A^+^ LE exhibits a hypoxic phenotype. **(a)** Boxplots comparing the expression levels of functional markers between different types of epithelial cells. **(b)** Violin plots comparing the expression level of HIF-1α in eIF1A^+^ LE between different types of tissues (PCT, PCa, low-risk PCa, high-risk PCa, LgPAC, HgPAC, IDC and DAC). **(c)** GSEA analysis of *EIF1A^hi^* versus *EIF1A^lo^* epithelial cells in the scRNA-seq validation cohort based on BIOCARTA, PID and HALLMARK database. **(d)** Correlation analysis between the AUCell score of HIF-1 TF pathway and EIF pathway in the scRNA-seq validation cohort. **(e)** Representative IMC images showing the staining of panCK, CK8, eIF1A and HIF-1α in different types of PCa. **(f)** The EIF and HIF-1 TF pathway AUCell score of malignant epithelia on ST slides. **(g)** Representative mIF staining images showing the co-localization of CK8, eIF1A and HIF-1α in different types of PCa (left); bar plots comparing the mean gray value of HIF-1α staining and the proportion of HIF-1α^+^ eIF1A^+^ LE in eIF1A^+^ LE between different types of PCa (right).

At a transcriptomic level, *EIF1A* expression was positively correlated with *HIF1A* in TCGA database (**Supplementary Figure 3a**). To better validate the hypoxic signature of eIF1A^+^ LE, we utilized a “single-cell RNA-seq (scRNA-seq) validation cohort” comprising 14 PAC samples (**Supplementary Table 2**). We found that classical carcinogenic signaling pathways (MYC targets V1, P53 pathway, TGF-β) and PCa-specific pathways (AR pathway, Androgen response) were enriched in *EIF1A*^hi^ epithelia. Notably, protein translation (EIF pathway) and hypoxia response pathways (HIF1 TF pathway, Hypoxia, Angiogenesis) were also activated in *EIF1A*^hi^ epithelia (**Figure 3c** and **Supplementary Figure 3b**). *EIF1A*, an important member of eukaryotic initiation factor (EIF), was significantly positively correlated with the AUCell score of EIF pathway in TCGA database, indicating high expression of eIF1A represented the activation of the EIF process and protein translation (**Supplementary Figure 3c**). Further, the activation level of EIF pathway in epithelia was significantly positively correlated with that of HIF-1 TF pathway, both at single-cell and bulk transcriptomic levels, reflecting a potential regulatory relationship between EIF induced protein translation and HIF-1α mediated hypoxia response (**Figure 3d** and **Supplementary Figure 3d-e**).

Spatially, we exhibited the greater HIF-1α expression in eIF1A^+^ LE in high-risk PCa subtypes (**Figure 3e**). We also performed a validation in our in-house spatial transcriptomic sequencing (ST-seq) data ^19^, confirming that EIF^hi^ epithelia also had higher activation levels of HIF-1 TF pathway, with a significantly positive correlation (**Figure 3f** and **Supplementary Figures 3f-h**). mIF staining verified greater mean gray levels of HIF-1α and higher HIF-1α^+^/eIF1A^+^ LE abundance in HgPAC, IDC and DAC compared to LgPAC (**Figure 3g** and **Supplementary Figure 4a**). Paired tests using each sample as a unit (e.g., samples containing both IDC and LgPAC components were included in paired comparisons between IDC and LgPAC) yielded consistent results **(Supplementary Figure 4b)**.

### eIF1A^+^ LE-centered “cold” and hypoxic immune microenvironment

When comparing the expression level of functional markers among epithelial sub types, we also found a higher expression of PD-L1 in eIF1A^+^ LE. IDO-1, which is the key enzyme of tryptophan metabolism, a downstream protein of HIF-1α, and a factor mediated hypoxia induced immunosuppression, was also overexpressed in eIF1A^+^ LE (**Figure 3a**). Therefore, we further investigated the immune microenvironment heterogeneity surrounding eIF1A^+^ LE that may be associated with PCa progression.

We first identified several cell types that correlate with clinical features (**Figure 4a**). Antitumor immune cells PD-1^-^ T cells, CD163^-^ Macrophages, and other APCs were associated with negative bladder neck invasion and were favorable for OS and PFI (**Figure 4a**, **Box 1-3**). PD-1^+^ T cells were enriched in patients with high tumor quantification and high Stage (**Figure 4a**, **Box 4**). We also found that stromal cells including myofibroblasts and SMC were negatively related with progression (**Figure 4a**, **Box 5-6**). Compared with DAC, other APCs were enriched in both LgPAC and IDC, which may indicate that the malignancy of DAC was associated with antigen presentation deficiency (**Figure 4a**, **Box 3**). Overall, these data revealed novel associations between TME cells and clinical outcomes, especially for immune cells.

**Figure 4.**
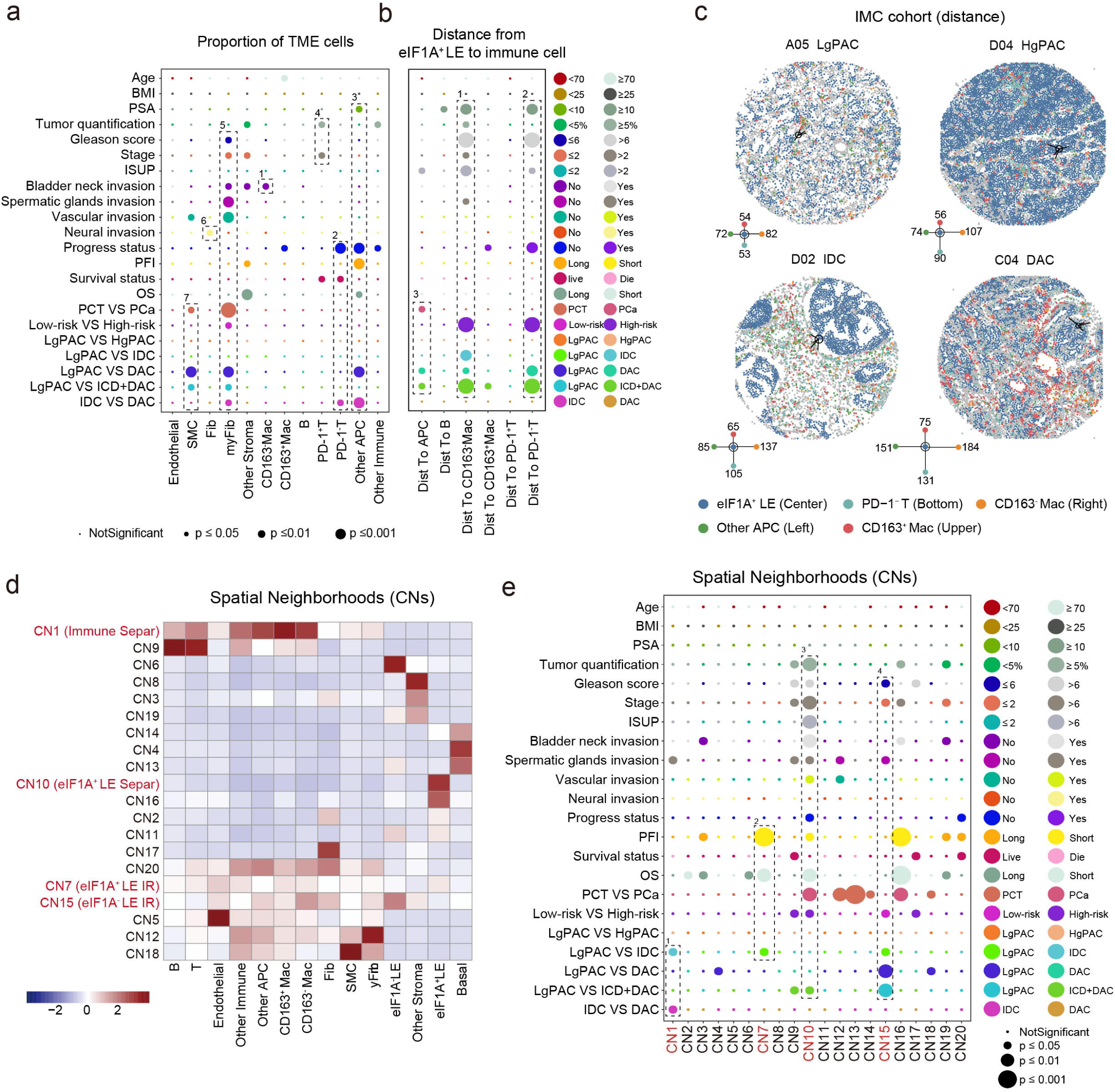
eIF1A^+^ LE-centered “cold” immune microenvironment. **(a)** Dot plot showing the association between the proportion of TME cells with clinical features. **(b)** Dot plot showing the association between the distance from eIF1A^+^ LE to immune cells with clinical features. **(c)** Representative IMC images showing the distance from eIF1A^+^ LE to PD-1^-^ T cells, CD163^-^ macrophages, CD163^+^ macrophages and other APC in different types of PCa. **(d)** Heatmap showing the components of 20 identified CNs. **(e)** Dot plot showing the association between the proportion of CNs with clinical features.

Subsequently, we assessed the cellular interactions in different subtypes through interaction analysis (**Supplementary Figure 5a**). We found that the avoidance between immune cells and eIF1A^+^ LE was significantly enhanced in high-risk compared to low-risk PCa (**Supplementary Figure 5a**, **Box 1**). Therefore, we further evaluated the distances between different immune cell types and eIF1A^+^ LE, and assessed the association of the distances with different clinical features (**Figure 4b**). The results revealed that for antitumor cells such as PD-1^-^ T cells, CD163^-^ Macrophages, and other APCs, their distances from eIF1A^+^ LE were positively correlated with malignant features including higher PSA, greater pathologic grade, progressed stage, malignant histologic types (IDC and DAC, **Figure 4b**, **Box 1-3** and **Supplementary Figure 5b**). Subsequently, we demonstrated the distance of eIF1A+ LE to key immune cells in different histological subtypes by cell spatial distribution mapping. It was found that HgPAC exhibited relatively sparse immune cell infiltration; in IDC, eIF1A^+^ LE were separated by a BE-stromal barrier, while in DAC, immune cells failed to infiltrate into eIF1A^+^ LE regions. Collectively, these advanced subtypes all developed “cold” immune microenvironments centered around eIF1A^+^ LE populations. (**Figure 4c**).

The results of cellular neighborhoods (CNs) also suggested consistent immune landscapes. We defined a total of 20 CNs by identifying the 10 nearest spatial neighbors for each cell (**Figure 4d** and **Supplementary Figure 5c**). CN10, named eIF1A^+^ LE separation CN, was abundant in tumors, especially high-risk PCa, and it was associated with multiple malignant pathological features and poor prognosis (**Figure 4e**, **Box 3, Supplementary Figure 5d-e**). CN7, which consisted of immune cells and eIF1A^+^ LE, represented an eIF1A^+^ LE immune response CN and was less enriched in IDC compared to LgPAC. CN7 was also associated with unfavorable prognosis, suggesting the absence of tumor-killing CNs in high-risk PCa (**Figure 4e**, **Box 2, Supplementary Figure 5d-e**). CN15 was an eIF1A^-^ LE immune response CN which was enriched in LgPAC and significantly reduced in IDC and DAC, implying that the less malignant potential of LgPAC might be attributed to effective immune recognition and elimination of eIF1A^-^ LE (**Figure 4e**, **Box 4** and **Supplementary Figure 5d-e**). Interestingly, as a neighborhood composed of T, CD163^-^ Mac, and other immune cells, the immune separation CN1 was highly enriched in IDC, suggesting that IDC has an immune exclusion feature (**Figure 4e**, **Box 1** and **Supplementary Figure 5d-e**).

Given that eIF1A^+^ LE exhibited a hypoxic phenotype, we further quired whether the aforementioned “cold” TME existed in a hypoxic niche through HIF-1α^+^ patch analysis. By assigning cells with high HIF-1α expression as hub cells, contiguous clusters of at least 10 HIF-1α^+^ cells with cells within their surrounding 10μm-radius microenvironment were designated as HIF-1α^+^ patches (**Figure 5a**). In agreement with the hypothesis, eIF1A^+^ LE was significantly more enriched inside than outside HIF-1α^+^ patches (**Figure 5b**). The HIF-1α^+^ patches also showed fewer PD-1^-^ T cells, CD163^-^ Macrophages and CD163^+^ Macrophages while more PD-1^+^ T cells, complying with the cellular characteristics of the above “cold” TME (**Figure 5b**). These findings demonstrate that across different high-risk subtypes of PCa, a consistent eIF1A^+^ LE-centered “cold” and hypoxic immune microenvironment emerged, which cooperates with eIF1A+ LE to drive disease progression.

**Figure 5.**
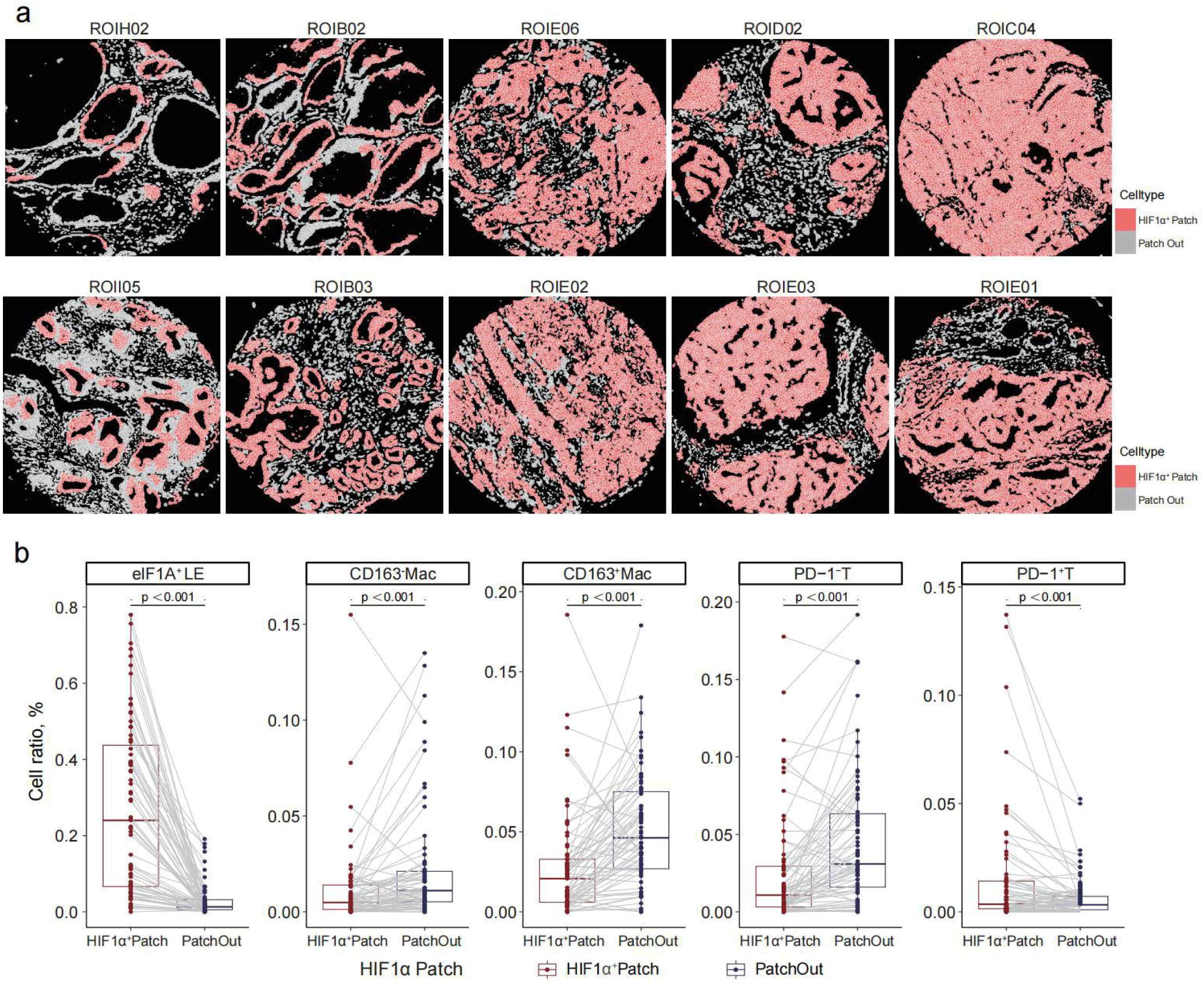
“Cold” immune microenvironment exists in a hypoxic niche. **(a)** Representative IMC images showing the location of HIF-1α^+^ patch in PCT and different types of PCa. **(b)** Boxplots comparing the proportion of eIF1A^+^ LE, PD-1^-^ T cells, PD-1^+^ T cells, CD163^-^ macrophages and CD163^+^ macrophages between inside and outside HIF-1α^+^ patches.

### HHT suppresses PCa growth by alleviating hypoxia and enhancing immune response

Our previous study discovered that protein translation was upregulated in small cell-like PCa and promoted disease progression, and translation inhibitor HHT was demonstrated to exert anti-cancer efficacy in PCa ^20^. The eIF1A and its associated EIF pathway identified in the current study were critical regulators of protein translation, whose downstream process could also be inhibited by HHT. Thus, we further investigated whether the treatment efficacy of HHT in PCa involved hypoxia alleviation and immune response enhancement.

Similar to previous findings, the growth of RM1 subcutaneous tumors could be significantly inhibited by HHT (**Figure 6a-b**). RNA-seq analysis of subcutaneous tumors suggested HHT treatment induced differentially expressed genes were associated with the alteration of response to hypoxia, positive regulation of T cell activation, myeloid cell activation involved in immune response, and tumor epithelial inhibitory pathway (negative regulation of epithelial cell proliferation, migration and mesenchymal transition) (**Figure 6c**). mIF staining revealed a significant decrease in HIF1α^+^ eIF1A^+^ LE in the tumors after HHT treatment (**Figure 6d**). CD163^-^ macrophages and CD8^+^ T cells were significantly increased after HHT treatment (**Figure 6e-f**).

**Figure 6.**
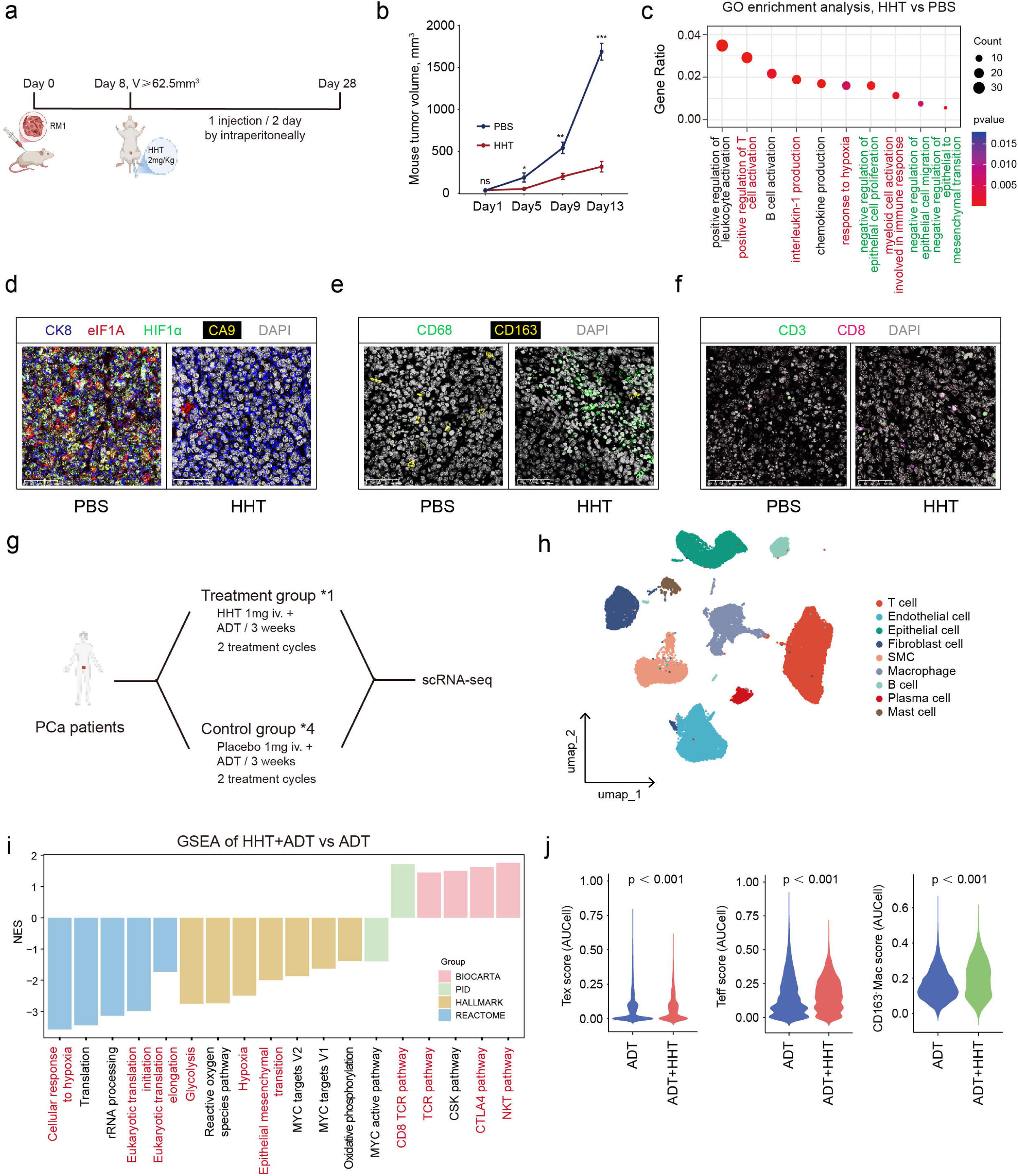
HHT suppresses PCa growth by alleviating hypoxia and enhancing immune response. **(a)** Diagram of mouse model construction strategy for HHT treatment. **(b)** The mouse tumor volum comparisons between HHT and control group. **(c)** GO enrichment analysis for the differentially expressed genes between HHT and control group based on RNA-seq. **(d-f)** Representative mIF staining images showing the distribution of eIF1A^+^ LE (CK8, eIF1A; d), hypoxic niche (HIF-1α, CA9; d), macrophages (CD68 CD163; e) and T cells (CD3, CD8; f) in HHT and control groups. **(g)** Diagram of the investigator-initiated study of HHT + ADT therapy (Treatment group) versus Placebo + ADT (Control group) in PCa patients and the subsequent scRNA-seq analysis. **(h)** UMAP plot of single cells from the scRNA-seq of 1 patient underwent HHT + ADT therapy and 4 patients underwent placebo + ADT therapy. **(i)** GSEA analysis of epithelial cells from Treatment group versus Control group based on BIOCARTA, PID, HALLMARK and REACTOME database. **(j)** Violin plots comparing the AUCell score of Exhausted T cells (Tex; left), effector T cell (Teff; middle) and CD163^-^ macrophage (right) signatures between Treatment group versus Control group.

Furthermore, we carried out an investigator-initiated study and investigated the molecular characteristics with scRNA-seq (**Figure 6g-h**). Compared with using ADT alone, HHT+ADT effectively inhibited malignant pathways such as protein translation, hypoxia, and epithelial mesenchymal transition, and enhanced immune-related pathways such as TCR, CTLA4, and NKT (**Figure 6i**). Moreover, there was a significant decrease in exhausted T cell score and a dramatic activation of anti-cancer immune process after adding HHT, represented by the effector T cell score and CD163^-^ macrophage score (**Figure 6j**). These results demonstrated that HHT treatment could alleviate hypoxia and enhance immune response in PCa, thus exerting anti-cancer efficacy, which further validated the function of eIF1A^+^ LE and the “cold” TME in advanced PCa.

## Discussion

Previous research on PCa has encountered significant difficulties in accurately delineating the spatial distribution and interactions among various pathological subtypes ^21^. This study represents a pioneering application of IMC technology to analyze the spatial microenvironmental characteristics of PCa and its subtypes. It identifies a hypoxia-responsive eIF1A+ tumor epithelial cell that is significantly correlated with advanced disease stages and a poorer prognosis. Surrounding these eIF1A+ tumor cells and hypoxic niches, different pathological subtypes exhibit ’cold’ tumor characteristics, thereby revealing the progression pattern of PCa. This investigation not only clarifies the relationship between the protein translation process, as indicated by the targeting of eIF1A, and hypoxia, along with its therapeutic implications for PCa, but also highlights the unique advantages of spatial proteomics technology. This study provides a novel perspective for the development of precision treatment strategies in PCa.

eIF1A is a eukaryotic translation initiation factor that is integral to the process of protein translation ^22^. In the realm of pan-cancer research, mutations or aberrant expression of its encoding gene, *EIF1A*, have been closely linked to the initiation, progression, and prognosis of various malignancies ^23,24^. Nevertheless, its specific role in PCa remains inadequately defined. This study aims to fill this knowledge gap by being the first to characterize eIF1A+ epithelial cells in the context of PCa and to examine their behavior under hypoxic conditions. Our findings indicate that eIF1A+ epithelial cells exhibit enhanced protein translation activity in hypoxic environments, potentially shedding light on the relationship between protein translation and hypoxia in PCa. Employing IMC technology, we observed that eIF1A+ tumor epithelial cells display elevated expression levels of HIF1 α and VEGFA. Single-cell analysis revealed that tumor epithelial cells with high *EIF1A* expression show significantly increased activity in the ’hypoxia response’ and ’HIF-1 signaling pathway.’ Furthermore, spatial transcriptomic analysis demonstrated a significant co-localization between heightened protein translation and increased activity of the HIF pathway, a correlation that was consistently supported by bulk RNA-seq data. Additionally, through tissue section staining, we further corroborated the relationship in localization and expression between eIF1A, which indicates protein translation, and HIF-1α, a recognized marker of hypoxia.

Our research corroborates previous investigations, highlighting considerable variability in clinical survival durations among distinct pathological subtypes of PCa^21^. This variability may be linked to differences in the tumor microenvironment associated with these subtypes.

There has been limited examination of the immune microenvironment in PCa across various clinical classifications, although variations in immune infiltration have been noted in localized PCa. In contrast, metastatic PCa and CRPC are typically regarded as existing within an immunosuppressive environment when compared to localized PCa ^25–27^. Nonetheless, prior studies have been insufficient in delineating the immune microenvironment characteristics of the various pathological subtypes of PCa. Our findings indicate that different pathological subtypes of PCa share a common trait of exhibiting a ’cold’ tumor phenotype, albeit to differing extents. In subtypes characterized by higher malignancy, as compared to PCT and Lg-PAC, the tumor microenvironment displays a more pronounced immunosuppressive effect surrounding eIF1A+ epithelial cells and hypoxic niches.

Previous investigations conducted by our research team have established a significant correlation between the aggressive phenotype of SCLPC and the activation of protein translation mechanisms ^28^. HHT, the sole protein translation inhibitor currently approved for clinical use, has exhibited antitumor efficacy across various malignancies ^29^. Our prior studies have demonstrated that HHT effectively hinders the progression of CRPC ^28^. Building upon these findings, which elucidate the role of protein translation in the development of hypoxic and immune ’cold’ tumor microenvironments, and bolstered by comprehensive animal studies, we have launched an investigator-initiated clinical trial. Utilizing single-cell transcriptome sequencing, we confirmed that, in comparison to ADT alone, the pathways associated with protein translation and hypoxia in the epithelial cells of patients were markedly downregulated following treatment with HHT in conjunction with ADT. Additionally, we noted a significant increase in the proportion of T cells, a considerable enhancement in the cytotoxic T cell score, and a notable rise in the scores of the M1 pro-inflammatory and anti-tumor macrophage subtypes. These observations suggest that HHT combination therapy may partially rectify the ’cold’ immune microenvironment characteristic of PCa, thereby facilitating the infiltration of tumoricidal immune cells into the tumor core and enhancing overall antitumor efficacy. This evidence indicates that HHT not only inhibits protein translation but also improves tumor hypoxia, thereby promoting the infiltration and functional activity of immune cells. Consequently, we posit that this discovery possesses substantial clinical significance, offering a novel strategy for the precision treatment of PCa and presenting opportunities for further translational research in this domain.

In the field of PCa research, prior investigations have predominantly relied on RNA sequencing or single-cell sequencing methodologies to examine various pathological subtypes ^30^. However, these techniques present notable limitations. While RNA sequencing offers extensive insights into gene expression profiles in both malignant and adjacent tissues, it does not elucidate the spatial distribution of cells within the tumor microenvironment or the interactions among them. Recent studies utilizing single-cell RNA sequencing have faced difficulties in detecting focally distributed rare subtypes, such as IDC and DAC ^31^. The focal distribution of these subtypes complicates the accurate identification of pertinent tumor and microenvironmental cells from a large cellular population, potentially resulting in unreliable findings. To overcome these challenges, the present study employs IMC technology to effectively construct an in situ single-cell protein atlas of various pathological subtypes of PCa. IMC technology facilitates the spatial characterization of protein expression levels at single-cell resolution, thereby enabling the precise identification of small regions of interest that encompass diverse pathological types ^32^. This methodology enhances the understanding of cellular functions and provides substantial support for elucidating the intricate tumor microenvironment associated with different pathological subtypes of PCa.

This study has some limitations. The tissue microarray technique utilized in this investigation facilitated an adequate sample size; however, it constrained our capacity to perform comprehensive comparisons across all tumor regions within individual patient samples. Additionally, this study has not yet delved into the specific molecular mechanisms through which eIF1A modulates protein translation, exacerbates hypoxia, and inhibits immune responses in tumor epithelial cells. The primary focus was to establish the causal relationship between eIF1A expression and malignant tumor phenotypes using fundamental in vivo and in vitro models, while also conducting preliminary assessments of its potential for clinical application.

In summary, this research has elucidated the common ’cold’ tumor characteristics across various pathological subtypes of PCa through IMC technology. It underscores the interplay between protein translation and hypoxia in PCa and their contributions to the development of an immunosuppressive microenvironment. HHT, a drug that targets protein translation, not only impedes tumor growth by inhibiting the protein translation process but also ameliorates hypoxia and enhances the immune microenvironment, thereby demonstrating considerable therapeutic promise. This study provides novel insights for the precision treatment of PCa and lays robust groundwork for future inquiries into the domains of protein translation and hypoxia, with the potential to stimulate further exploration into the regulatory mechanisms governing the PCa microenvironment.

## Methods

### Ethics Statement

The use of PCa tissues in this study was approved by the Clinical Research Ethics Committee of Zhongda Hospital Southeast University (Approval No.: 2024ZDSYLL382-P01). Written informed consent was obtained from all patients. Additionally, we conducted an investigator-initiated clinical trial, which included one advanced patient who received androgen deprivation therapy (ADT) combined with HHT treatment, as well as four patients who received only ADT treatment preoperatively. The study design and informed consent documents for research participants complied with relevant regulations and ethical principles and were approved by the Ethics Committee of the First People’s Hospital of Nantong (Approval No.: 2025KT076).

### Human specimens and clinical data collection

The 71 cores on the TMA were collected from 47 clinical samples from patients diagnosed with PCa from 2019-10 to 2023-09, with a follow-up until 2025-04. All samples were diagnosed by certified pathologists following surgical resection. The clinical data of the patients included age, gender, prostate-specific antigen (PSA) levels, and Gleason score (GS), while the pathological data encompassed tumor size, grade, presence of lymph node metastasis, and additional relevant information. All data were extracted from the electronic medical record system and subsequently reviewed and confirmed by professional pathologists to ensure accuracy and completeness. Detailed clinical information for all enrolled patients is provided in (**Supplementary Table 1**). Clinical samples were utilized for the preparation of tissue microarrays (TMA) and multiplex immunohistochemistry paraffin sections.

### Tissue Microarray (TMA) Preparation

TMA is constructed by selecting representative cores with a diameter of 1 mm from surgical tumor specimens. The specific sampling locations were determined according to the research requirements. The specific steps for TMA construction are as follows: the tumor core area within the postoperatively resected prostate tumor specimens, adjacent normal tissue distant from the tumor, or other tissue samples conforming to the research objectives are processed into FFPE blocks. A representative core is taken from each tissue area FFPE block (with a diameter of 1 mm) and arranged into a TMA that represents 46 patients from 77 blocks.

### Sample Collection and Sequencing

To thoroughly analyze the cellular heterogeneity of PCa tissues, we performed single-cell RNA sequencing (scRNA-seq) on tumor samples from several PCa patients’ post-surgery and integrated these with scRNA-seq samples previously generated by our research group. Additionally, in our ongoing clinical trials, we conducted scRNA-seq on the resected tissues of patients after their treatment concluded. The specific procedures are as follows: fresh tumor tissue samples were washed with Hanks’ Balanced Salt Solution (HBSS) at 4°C, followed by digestion with collagenase to prepare a single-cell suspension. Undigested tissue fragments and cell clusters were removed by centrifugation, and the single-cell suspension was collected. Single-cell capture was performed using the 10x Genomics Chromium platform, with each cell encapsulated in individual microdroplets. Subsequently, in accordance with the standard operating procedures of 10x Genomics, single cells underwent reverse transcription, cDNA amplification, and library construction. The constructed libraries were sequenced on the Illumina NovaSeq 6000 platform using a 150 bp paired-end sequencing (PE150) strategy. Upon completion of sequencing, data processing was conducted using the Cell Ranger software, which included quality control, alignment to the human reference genome (GRCh38), and generation of the gene expression matrix.

### Imaging Mass Cytometry (IMC) Preparation Process and Image Computation

#### IMC Panel Design and Antibody Selection

The selection of the IMC staining panel is based on an analysis of target cell types and their biological characteristics. Antibodies undergo rigorous screening and validation, including dual verification through immunofluorescence (IF) and IMC, to ensure their specificity and signal intensity. The panel encompasses a variety of cell types and markers, including immune cells (such as T cells, B cells, and myeloid cells) as well as tumor cell-related markers. Additionally, it includes markers associated with biological and metabolic processes, such as hypoxia and energy metabolism. The antibody dilution is optimized through a series of dilution experiments to achieve the optimal staining effect. Comprehensive antibody information is detailed in the **Supplementary Table 3**.

#### IMC Sample Preparation

Samples were obtained from tissue microarrays (TMA) and serially sectioned to a thickness of 5 μm. The sectioning process proceeded as follows: Sections were dewaxed at 70°C, followed by antigen retrieval at 95°C. Non-specific binding sites were blocked using Dako’s protein-free blocking solution and incubated for 45 minutes. A mixture of metal-labeled antibodies, diluted in 1% BSA/PBS, was applied to the sections and incubated overnight at 4°C. The sections were then washed with 0.2% Triton X-100 and 1× PBS, followed by nuclear counterstaining, which was performed at room temperature for 30 minutes. Subsequently, the sections were washed with ultrapure water and air-dried naturally. IMC images were acquired using the Hyperion imaging system.

#### IMC Antibody Labeling and Validation

The antibody information is detailed in **Supplementary Table 3**. All antibodies were optimized and validated across various tissues, including the spleen, tonsil, appendix, placenta, thymus, normal lung tissue, and PCa tissue, to ensure their specificity and signal strength.

#### IMC Imaging and Data Acquisition

Following the acquisition of IMC images, each TMA core is individually imaged using the imaging system. The tissue is scanned through laser ablation, where the ablated tissue plume is ionized by a plasma source. Signals from metal-tagged antibodies are then detected via time-of-flight mass spectrometry (TOF-MS). Ultimately, images corresponding to each antibody are reconstructed based on the metal abundance of each pixel. The staining results are reviewed by professional pathologists.

IMC image preprocessing and cell segmentation involve a comprehensive data analysis pipeline that includes spillover signal compensation, image denoising, image contrast enhancement, and cell segmentation. The spillover signals in each channel are corrected using a previously defined spillover matrix ^33^. Median filtering is applied to suppress noise, followed by intensity adjustment using the Matlab function imadjust to optimize the intensity distribution. Finally, connectivity-aware segmentation methods are employed to accurately segment individual cells or components in the IMC images ^34^.

#### Image Processing and Spatial Information Analysis

Overflow signals are filtered out, and median filtering is applied to reduce noise in the signals from each channel^33^. The contrast between the signals and the background is enhanced through linear adjustment.

During cell segmentation, membrane protein signals located more than 20 μm away from the nucleus center are excluded. Marker expression data undergo hyperbolic sine transformation and are subsequently normalized using the min-max method. Batch effects are corrected using the R software Harmony (version 0.1.0)^35^, and cells are clustered using Rphenograph (version 0.99.1, k=100)^36^. The clustering results are employed for cell type (CT) annotation. The cellular neighborhood (CN) of each cell consists of its 10 nearest neighboring cells. The neighborhoods are clustered and annotated using the k-means clustering method (k=15) and validated through Voronoi diagrams. Permutation tests are conducted using imcRtools (version 1.5.4) to analyze the interactions between cell types within the selected region of interest (ROI). Differences in interactions between various groups are compared using Student’s t-test, with P<0.05 indicating significant spatial interactions between cells across different groups.

### scRNA-seq Data Processing and Analysis

Data Preprocessing and Dimensionality Reduction: AnalysisInitially, the data were normalized using the Seurat package (v.4.0.3) with the LogNormalize method (scale.factor = 10000). Subsequently, the top 2000 highly variable feature genes were identified using the FindVariableFeatures function with the vst method. Following this, all genes were standardized using the ScaleData function, and PCA dimensionality reduction was performed with the RunPCA function.

#### Batch Effect Correction and Cell Clustering

To address potential batch effects, the Harmony package (v1.0) was utilized for dimensionality reduction and integration. The number of principal components was determined using the Elbow plot, with components accounting for over 90% of the cumulative variance selected for subsequent analysis. Based on the dimensionality reduction results from Harmony, cell clustering analysis was conducted using Seurat’s FindNeighbors and FindClusters functions, evaluating clustering effects at various resolutions (0.05, 0.1, 0.2, 0.3, 0.5, 0.8, 1.0). An appropriate resolution was ultimately chosen for UMAP dimensionality reduction visualization (RunUMAP), and cell types were annotated according to the clustering results.

#### Cell subpopulation analysis

We specifically focused on the epithelial cell subpopulations, which were categorized into *EIF1A*-High and *EIF1A*-Low subgroups based on the median expression level of the *EIF1A*.

#### Gene Set Scoring

The AUCell algorithm, along with the AUCell package (v1.16.0), was employed to calculate the transcriptional signature scores for individual cells and samples.

#### CellChat Analysis

To assess the intercellular communication network, single-cell data were analyzed using CellChat (v1.1.0). Initially, the single-cell patients were categorized into high-expression and low-expression groups based on their Gleason scores. Following this, CellChat objects were constructed for each group, and ligand-receptor interaction analyses were conducted. The communication probabilities between cells were calculated using the computeCommunProb and aggregateNet functions of CellChat, and communication network diagrams were generated. Furthermore, a comparative analysis of the communication networks between different groups was performed to elucidate differences in intercellular communication.

#### Identification of Differentially Expressed Genes and Functional Enrichment Analysis

To investigate the potential mechanisms underlying the high and low expression of *EIF1A* in epithelial cell subpopulations, as well as the effects of combination therapy with HHT and ADT compared to ADT alone, the FindMarkers function in Seurat was utilized to identify differentially expressed genes, applying a logFC threshold of 0 and a min.pct of 0.005. Following this, analyses were performed using clusterProfiler (v4.4.0) and GSEA (v1.42.0) based on the differentially expressed genes, which included the GO, KEGG, BIOCARTA, REACTOME, and PID databases.

### Processing and Analysis of Spatial Transcriptomics Data

#### Data Preprocessing

We utilized the Seurat package to analyze published spatial transcriptomics data of PCa. Initially, each spot was normalized using the SCTransform method to correct for technical variations and stabilize the data. Subsequently, the PCA dimensionality reduction technique was employed to reduce data dimensions through the RunPCA function. Based on the PCA results, the number of principal components for subsequent analysis was determined using the ElbowPlot method, followed by data clustering through the FindNeighbors and FindClusters functions. This process ensures that the dimensionality reduction and clustering analyses accurately reflect the spatial heterogeneity within the tissue.

#### Epithelial Cell Annotation

To further analyze the characteristics of epithelial cells in spatial transcriptomic data, we utilized the previously integrated single-cell data from our hospital cohort, employing the CARD (v.1.1) deconvolution method in conjunction with known gene markers to annotate epithelial cells within the spatial transcriptomic data.

#### Visualization of Gene Expression

To intuitively demonstrate the expression levels and spatial distribution of specific genes, such as *EIF1A* and *HIF1A*, we utilized the SpatialFeaturePlot function to generate spatial distribution maps of gene expression.

#### Gene Set Scoring and Correlation Analysis

To evaluate the gene set expression characteristics of each spot, we utilized the GSVA package (v.1.42.0) to perform scoring analysis on the gene expression matrix corresponding to each spot. Subsequently, we conducted correlation analysis to assess the relationships between different gene set scores, visualizing the results to reveal the spatial cooperative patterns of these critical biological processes.

### Mouse bulk RNA-seq data processing and analysis

We used the “DESeq2” package to identify differentially expressed genes (DEGs) based on the criteria of p-adj < 0.05 and |logFC| > 1. Subsequently, we employed the “clusterProfiler” R package to conduct functional enrichment analysis of the differentially expressed genes, aiming to explore the biological functional differences between different groups. This included analysis based on the GO and KEGG databases.

## Statistical Analysis

In this study, all statistical analyses were conducted using R (v.4.0.5). Specifically, the Wilcoxon rank-sum test was employed to compare differences between two groups, which is suitable for non-normally distributed data. Simultaneously, the Kruskal-Wallis test was used to compare differences among multiple groups, also applicable to non-normally distributed data. Additionally, assuming that the data followed a normal distribution, the t-test was utilized to compare differences between two groups, while one-way analysis of variance (ANOVA) was applied to compare differences among multiple groups. For correlation analysis, we employed both Spearman and Pearson analyses to explore the relationships between the two factors, which are appropriate for non-normally distributed data. We utilized the survminer package (v.0.4.9.999) to conduct survival analysis via Kaplan-Meier curves, dividing individuals into high-expression and low-expression groups based on the determined optimal cutoff value. This study did not employ statistical methods to predetermine the sample size; however, the sample size threshold was referenced from previously published articles. Data collection and analysis were performed in a blinded manner, and no animals or data points were excluded.

### Multiplex Immunofluorescence (mIF) Staining

The mIF staining of target proteins in tissue sections was conducted using the Tyramide Signal Amplification (TSA) technique. The experimental procedure was as follows: First, paraffin-embedded tissue sections were baked at 65°C for 2 hours, followed by dewaxing and rehydration in xylene and graded ethanol solutions. Subsequently, antigen retrieval was performed in EDTA buffer at pH 8.0 or citrate buffer at pH 6.0, followed by washing with PBS. Endogenous peroxidase activity was inhibited using a 3% hydrogen peroxide solution at room temperature for 15 minutes. Next, non-immune goat serum was applied and incubated at room temperature for 30 minutes to block nonspecific binding. After removing the blocking solution, the primary antibody against the target protein was applied and incubated overnight at 4°C. Following PBST washing, an HRP-labeled secondary antibody was added and incubated at room temperature for 30 minutes. The TSA chromogenic reaction was then performed, with the specific duration determined based on preliminary experiments and prior experience. After washing with PBST, the aforementioned steps were repeated until all labeling was completed. Finally, the nuclei were counterstained with DAPI, the slides were mounted with antifade mounting medium, and images were observed and captured using a fluorescence microscope.

### Construction of Mouse Model and RNA-seq Sequencing

The experiment utilized 6 to 8-week-old male C57BL/6 mice, purchased from the Animal Experiment Center of Southeast University, and acclimatized for one week. The mice were housed under standard conditions, including a 12-hour light/dark cycle, with free access to food and water. All animal experiments were conducted at the Animal Experiment Center of Southeast University, strictly following the protocol approved by the Animal Research Committee of Southeast University, in compliance with the “Guide for the Care and Use of Laboratory Animals.” To construct the RM-1 PCa subcutaneous tumor model in mice, RM-1 cells in the logarithmic growth phase were digested, washed, centrifuged, and then diluted to a cell suspension of 1.0×10⁶ cells/ml. Using a 1 mL syringe, 200 µL of the cell suspension (containing 2×10⁵ RM-1 cells) was injected subcutaneously into the right dorsal region of the mice. After injection, the mice were randomly divided into an experimental group and a control group, with each group containing 4 mice. To validate the efficacy and mechanism of HHT treatment, the experimental group received intraperitoneal injections of HHT, while the control group received intraperitoneal injections of PBS. The treatment regimen was administered 3 times per week for 3 consecutive weeks. During the treatment period, we regularly monitored the body weight and tumor volume of the mice to assess treatment tolerance and preliminary efficacy. On day 3 following the last treatment, we collected tumor samples from the mice and conducted RNA-seq sequencing. The sequencing experiment was carried out, which generated high-quality transcriptomic data. Subsequently, we utilized software such as FastQC and HISAT2 to perform quality control, alignment, and quantification analysis on the raw data, resulting in a gene expression matrix for each sample.

## Acknowledgments

We are grateful to the staff of the Public Scientific Research Platform of Zhongda Hospital Southeast University, and Infinity Scope for technical assistance.

## Author contributions

Conceptualization: BX, YFC, SJ, MC

Methodology: YFC, LW, EH, CC, TW, SC, JZ

Investigation: LW, ZX, BZ, XD, RN

Visualization: LW, EH, YFC, CC, TW

Clinical samples collection: LW, ZX, BZ, XD

Project administration: BX, MC

Supervision: BX, DZ, SJ

Writing-original draft: LW, EH, YFC

Writing-review & editing: BX, YFC, WL, DZ, SJ, RN, MC

## Data and materials availability

All data are available in the main text or the supplementary materials.

## Ethics approval and consent to participate

All human tumor tissue samples were collected in accordance with the national and institutional ethical guidelines. The study design was approved by the Ethics Committee of Zhongda Hospital Southeast University. The approval ID is 2024ZDSYLL382-P01.

## Consent for publication

All authors approved the manuscript for submission and gave their consent for publication.

## References

1. Siegel RL, Giaquinto AN, Jemal A. Cancer statistics, 2024. CA Cancer J Clin. 2024;74(1):12–49.

2. Watson PA, Arora VK, Sawyers CL. Emerging mechanisms of resistance to androgen receptor inhibitors in prostate cancer. Nat Rev Cancer. 2015;15(12):701–711.

3. Loblaw DA, Virgo KS, Nam R, et al. Initial hormonal management of androgen-sensitive metastatic, recurrent, or progressive prostate cancer: 2006 update of an American Society of Clinical Oncology practice guideline. J Clin Oncol. 2007;25(12):1596–1605.

4. Kwon WA, Joung JY. Immunotherapy in Prostate Cancer: From a “Cold” Tumor to a “Hot” Prospect. Cancers (Basel). 2025;17(7).

5. Carretero R, Gil-Julio H, Vazquez-Alonso F, Garrido F, Castineiras J, Cozar JM. Involvement of HLA class I molecules in the immune escape of urologic tumors. Actas Urol Esp. 2014;38(3):192–199.

6. He X, Xu C. Immune checkpoint signaling and cancer immunotherapy. Cell Res. 2020;30(8):660–669.

7. Graff JN, Beer TM, Alumkal JJ, et al. A phase II single-arm study of pembrolizumab with enzalutamide in men with metastatic castration-resistant prostate cancer progressing on enzalutamide alone. Journal for immunotherapy of cancer. 2020;8(2).

8. Zhou M. Intraductal carcinoma of the prostate: the whole story. Pathology. 2013;45(6):533–539.

9. Tsuzuki T. Intraductal carcinoma of the prostate: a comprehensive and updated review. Int J Urol. 2015;22(2):140–145.

10. Porter LH, Lawrence MG, Ilic D, et al. Systematic Review Links the Prevalence of Intraductal Carcinoma of the Prostate to Prostate Cancer Risk Categories. Eur Urol. 2017;72(4):492–495.

11. Zhao T, Liao B, Yao J, et al. Is there any prognostic impact of intraductal carcinoma of prostate in initial diagnosed aggressively metastatic prostate cancer? Prostate. 2015;75(3):225–232.

12. Zhao J, Shen P, Sun G, et al. The prognostic implication of intraductal carcinoma of the prostate in metastatic castration-resistant prostate cancer and its potential predictive value in those treated with docetaxel or abiraterone as first-line therapy. Oncotarget. 2017;8(33):55374–55383.

13. Miura N, Mori K, Mostafaei H, et al. The Prognostic Impact of Intraductal Carcinoma of the Prostate: A Systematic Review and Meta-Analysis. J Urol. 2020;204(5):909–917.

14. Harkin T, Elhage O, Chandra A, et al. High ductal proportion predicts biochemical recurrence in prostatic ductal adenocarcinoma. BJU Int. 2019;124(6):907–909.

15. Ranasinghe W, Shapiro DD, Zhang M, et al. Optimizing the diagnosis and management of ductal prostate cancer. Nat Rev Urol. 2021;18(6):337–358.

16. Morgan TM, Welty CJ, Vakar-Lopez F, Lin DW, Wright JL. Ductal adenocarcinoma of the prostate: increased mortality risk and decreased serum prostate specific antigen. J Urol. 2010;184(6):2303–2307.

17. Ranasinghe WKB, Brooks NA, Elsheshtawi MA, et al. Patterns of metastases of prostatic ductal adenocarcinoma. Cancer. 2020;126(16):3667–3673.

18. Giesen C, Wang HA, Schapiro D, et al. Highly multiplexed imaging of tumor tissues with subcellular resolution by mass cytometry. Nat Methods. 2014;11(4):417–422.

19. Cheng Y, Liu B, Xin J, et al. Single-cell and spatial RNA sequencing identify divergent microenvironments and progression signatures in early-versus late-onset prostate cancer. Nat Aging. 2025.

20. Zou C, Li W, Zhang Y, et al. Identification of an anaplastic subtype of prostate cancer amenable to therapies targeting SP1 or translation elongation. Sci Adv. 2024;10(14):eadm7098.

21. Wasinger G, Oszwald A, Shariat SF, Compérat E. Histological patterns, subtypes and aspects of prostate cancer: different aspects, different outcomes. Current Opinion in Urology. 2022;32(6):643–648.

22. Chaudhuri J, Si K, Maitra U. Function of eukaryotic translation initiation factor 1A (eIF1A) (formerly called eIF-4C) in initiation of protein synthesis. Journal of Biological Chemistry. 1997;272(12):7883–7891.

23. Sehrawat U, Koning F, Ashkenazi S, Stelzer G, Leshkowitz D, Dikstein R. Cancer-Associated Eukaryotic Translation Initiation Factor 1A Mutants Impair Rps3 and Rps10 Binding and Enhance Scanning of Cell Cycle Genes. Molecular and Cellular Biology. 2019;39(3).

24. Krishnamoorthy GP, Davidson NR, Leach SD, et al. *EIF1AX* and *RAS* Mutations Cooperate to Drive Thyroid Tumorigenesis through ATF4 and c-MYC. Cancer Discovery. 2019;9(2):264–281.

25. Brea L, Yu J. Tumor-intrinsic regulators of the immune-cold microenvironment of prostate cancer. Trends in endocrinology and metabolism: TEM. 2025.

26. Novysedlak R, Guney M, Al Khouri M, et al. The Immune Microenvironment in Prostate Cancer: A Comprehensive Review. Oncology. 2024.

27. Wu W, Wang XA, Le W, et al. Immune microenvironment infiltration landscape and immune-related subtypes in prostate cancer. Frontiers in Immunology. 2023;13.

28. Zou C, Li WC, Zhang YZ, et al. Identification of an anaplastic subtype of prostate cancer amenable to therapies targeting SP1 or translation elongation. Science Advances. 2024;10(14).

29. Wang W, He L, Lin T, et al. Homoharringtonine: mechanisms, clinical applications and research progress. Frontiers in Oncology. 2025;15.

30. Zhao JE, Xu NW, Zhu S, et al. Genomic and Evolutionary Characterization of Concurrent Intraductal Carcinoma and Adenocarcinoma of the Prostate. Cancer Research. 2024;84(1):154–167.

31. Wong HY, Sheng QH, Hesterberg AB, et al. Single cell analysis of cribriform prostate cancer reveals cell intrinsic and tumor microenvironmental pathways of aggressive disease. Nature Communications. 2022;13(1).

32. Elhanani O, Ben-Uri R, Keren L. Spatial profiling technologies illuminate the tumor microenvironment. Cancer Cell. 2023;41(3):404–420.

33. Chevrier S, Crowell HL, Zanotelli VRT, Engler S, Robinson MD, Bodenmiller B. Compensation of Signal Spillover in Suspension and Imaging Mass Cytometry. Cell Systems. 2018;6(5):612-+.

34. Guadayol O, Thornton KL, Humphries S. Cell morphology governs directional control in swimming bacteria. Scientific Reports. 2017;7.

35. Korsunsky I, Millard N, Fan J, et al. Fast, sensitive and accurate integration of single-cell data with Harmony. Nature Methods. 2019;16(12):1289-+.

36. Levine JH, Simonds EF, Bendall SC, et al. Data-Driven Phenotypic Dissection of AML Reveals Progenitor-like Cells that Correlate with Prognosis. Cell. 2015;162(1):184–197.

