## Supplementary figure 1-5 for "A unified eIF1A^+^ luminal cells-centered hypoxic and “cold” tumor microenvironment promotes PCa progression among different subtypes"

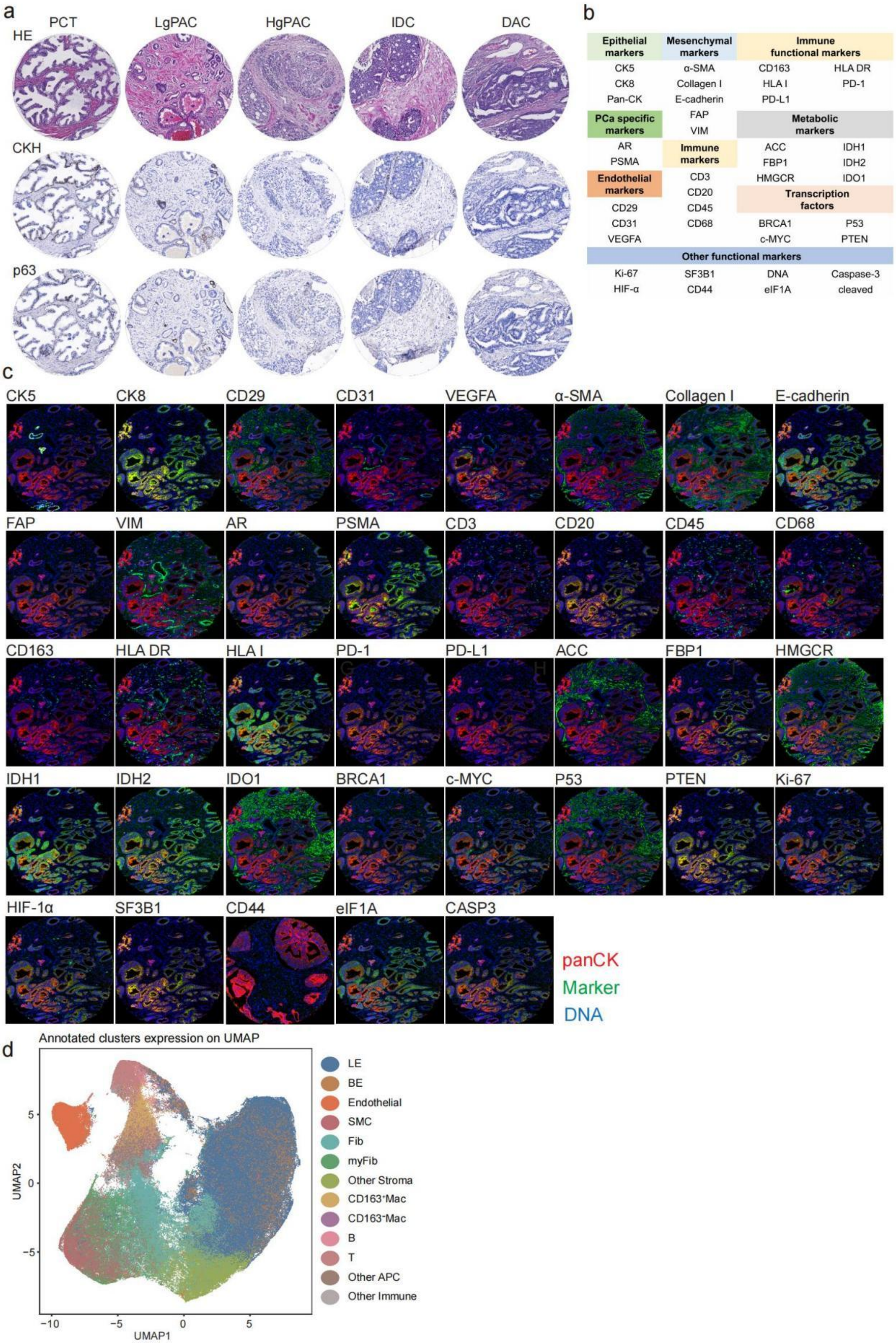


**Supplementary Figure 1. Preparation for IMC analysis**

(a) Hematoxylin & eosin, CKH, and p63 staining characteristics of tissue microarrays. (b) Target antibody in IMC analysis. (c) Fluorescence detection of target antibody alone. (d) UMAP plot of cell population distribution by clustering algorithm.

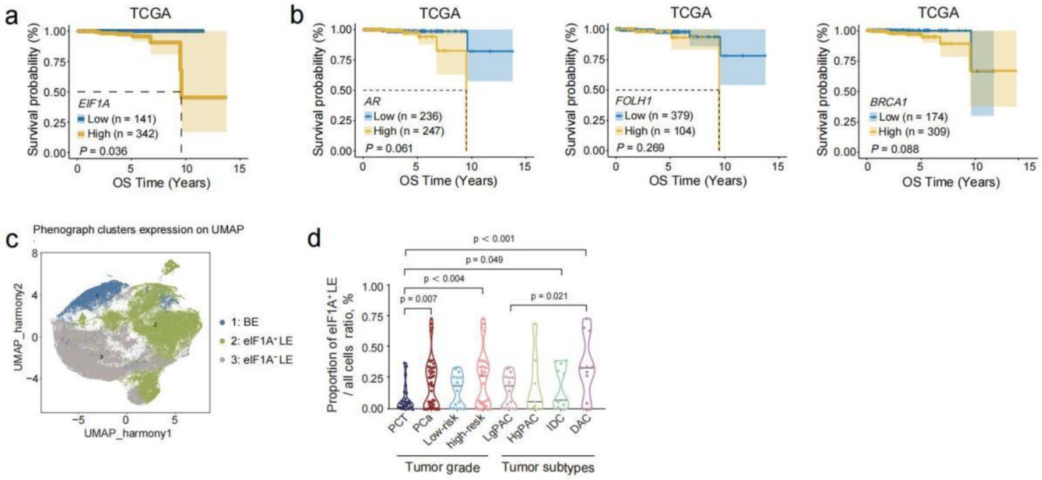


**Supplementary Figure 2. Prognostic analysis of classical oncogenes in TCGA, and proportional distribution of eIF1A$^+$LE in PCa.**

(a) The relationship between *EIF1A* expression levels and OS in the TCGA dataset. (b) The relationship between the expression levels of *AR, FOLH1*, and *BRCA1* and OS in the TCGA dataset. (C) UMAP plot showing the expression distribution of different phenotypic clusters (1: BE, 2: eIF1A$^+$LE, 3: eIF1A$^-$LE). (d) A violin plot of the proportions of eIF1A$^+$LE cells across different tumor grades and subtypes.

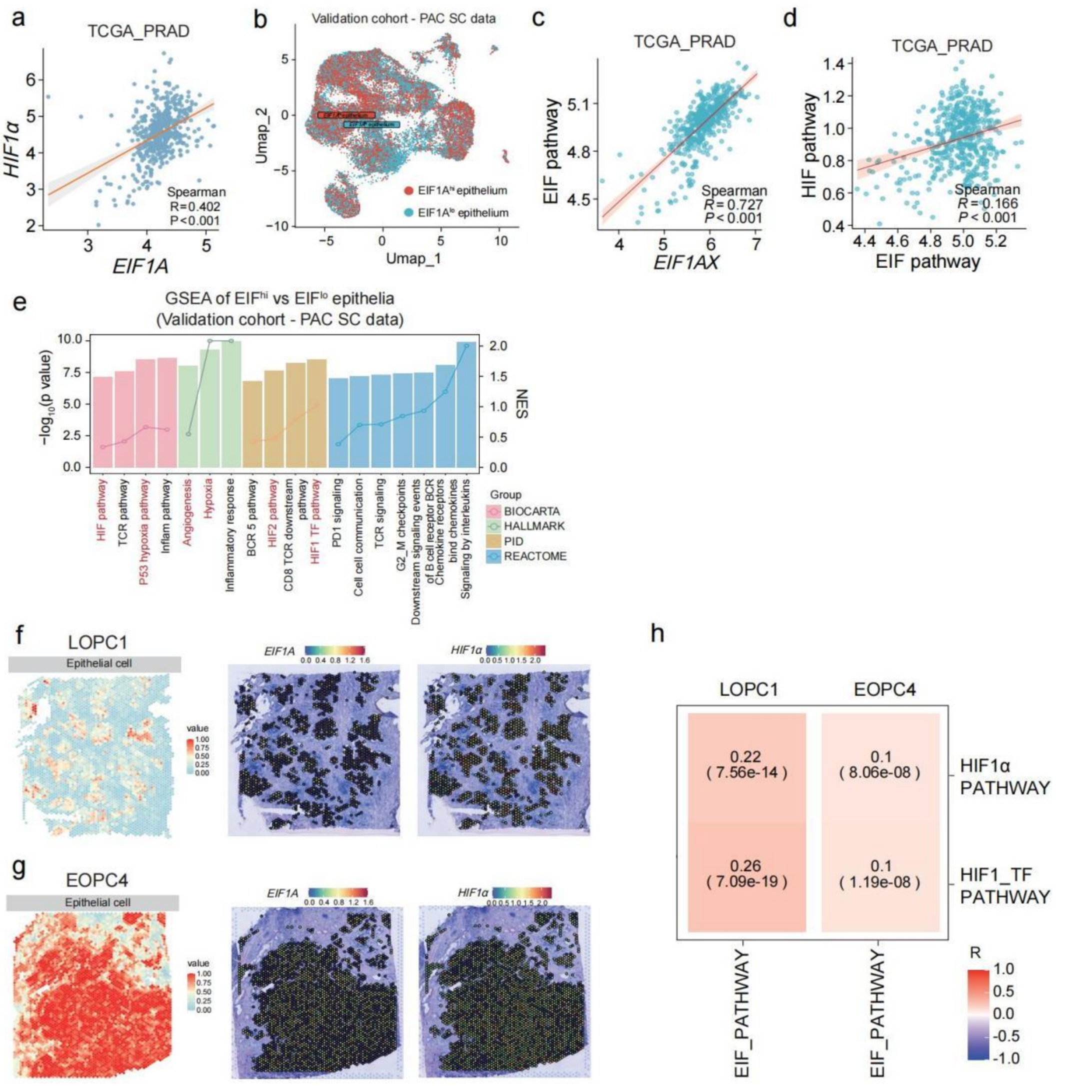


**Supplementary Figure 3. Correlation of expression patterns and signaling pathway activities of *EIF1A* and *HIF1α* in different cohorts.**

(a) Correlation of *EIF1A* with *HIF1α* in the TCGA-PRAD datase. (b) The UMAP plot of *EIF1A*$^{hi}$ and *EIF1A*$^{lo}$ epithelial cells in validation cohort - PAC SC data. (c) GSEA results for *EIF1A*$^{hi}$ and *EIF1A*$^{lo}$ samples in TCGA-PRAD. (d) Correlation of *EIF1A* with EIF pathway in the TCGA-PRAD datase. (e) Correlation of EIF pathway with HIF pathway in the TCGA-PRAD dataset. (f) GSEA results for *EIF* $^{hi}$ and *EIF* $^{lo}$ epithelium in the PAC SC data. (g-h) The expression of *EIF1A* and *HIF1α* in epithelial cells from spatial transcriptome samples LOPC1 and EOPC4. The left side shows the spatial distribution heatmap of epithelial cells, while the middle and right sides display the expression heatmaps of *EIF1A* and *HIF1a* in epithelial cells. (i) Correlation heatmap of EIF_PATHWAY with HIF1α_PATHWAY or HIF1_TF_PATHWAY in LOPC1 and EOPC4.

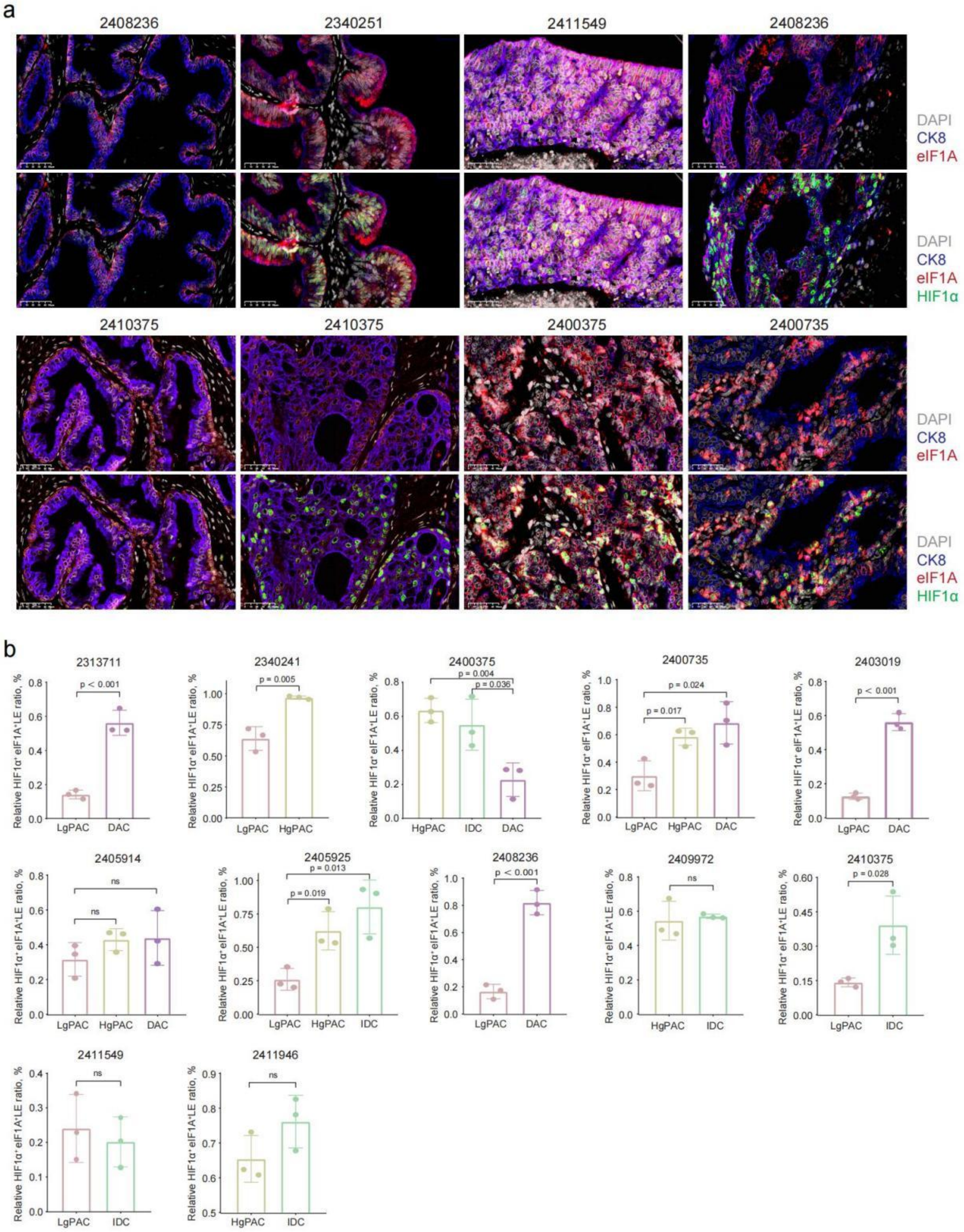


**Supplementary Figure 4. CK8-eIF1A-HIF1α multiplex immunofluorescence analysis in different tumor subtypes.**

(a) Multiplex immunofluorescence staining showing DAPI (blue), CK8 (red), eIF1A (yellow), and HIF1α (green) in different samples. (b) Histogram of $HIF1\alpha^{+}eIF1A^{+}LE$ ratios in different subtypes of a single sample.

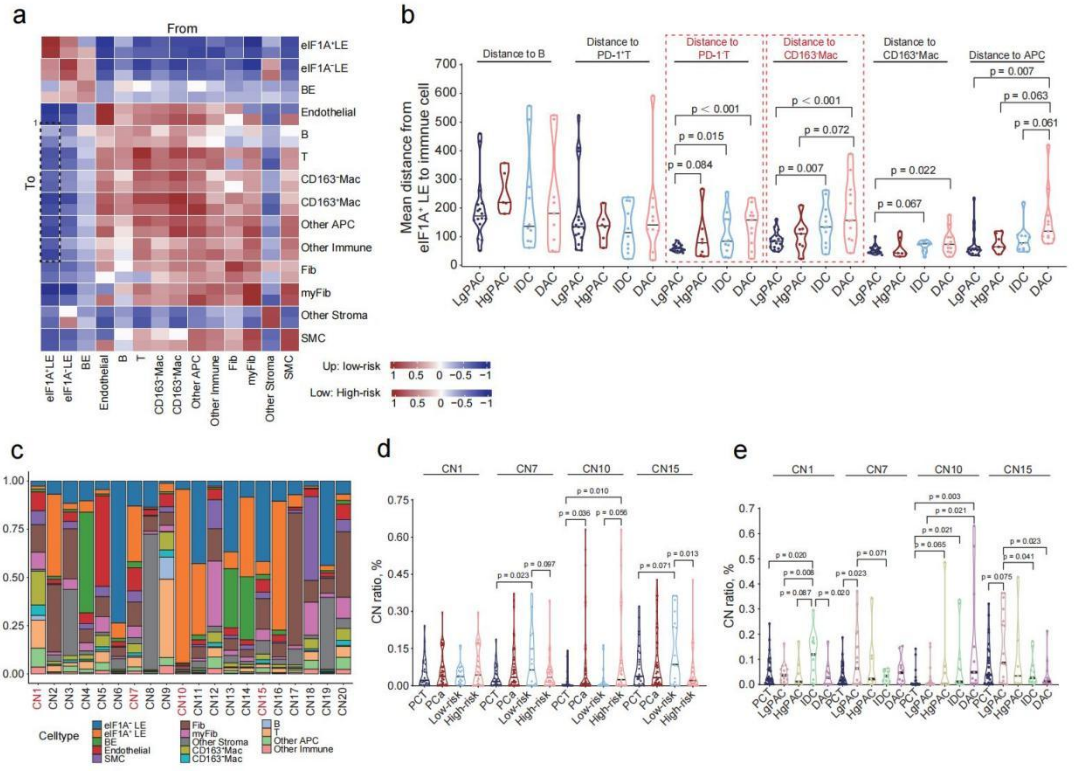


**Supplementary Figure 5. Cell spatial analysis centered on eIF1A⁺LE.**

Heatmaps depicting cell-cell interactions (red) or avoidance (blue) in low-risk and high-risk prostate cancer. The black boxes depict associations referenced in the text. (a) Violin plots showing the average distances from eIF1A⁺LE to different immune cell in different tumor subtypes. (c) Proportions of neighborhood assignments for each ROI. (d-e) Violin plots showing the proportions of four CNs (CN1, CN7, CN10, CN15) in different tumor grades and subtypes.
